# Tiny amphibious insects use tripod gait for seamless transition across land, water, and duckweed

**DOI:** 10.1101/2024.04.02.587757

**Authors:** Johnathan N. O’Neil, Kai Lauren Yung, Gaetano Difini, Holden Walker, M. Saad Bhamla

**Affiliations:** School of Chemical and Biomolecular Engineering, Georgia Institute of Technology, Atlanta, Georgia, United States

**Keywords:** *Microvelia*, water strider, semiaquatic, amphibious, duckweed, alternating tripod gait, neuston

## Abstract

Insects exhibit remarkable adaptability in their locomotive strategies across diverse environments, a crucial trait for foraging, survival, and predator avoidance. *Microvelia*, tiny 2-3 mm insects that adeptly walk on water surfaces, exemplify this adaptability by using the alternating tripod gait in both aquatic and terrestrial terrains. These insects commonly inhabit low-flow ponds and streams cluttered with natural debris like leaves, twigs, and duckweed. Using high-speed imaging and pose-estimation software, we analyze *Microvelia spp*.*’s* movement across water, sandpaper (simulating land), and varying duckweed densities (10%, 25%, and 50% coverage). Our results reveal *Microvelia* maintain consistent joint angles and strides of their upper and hind legs across all duckweed coverages, mirroring those seen on sandpaper. *Microvelia* adjust the stride length of their middle legs based on the amount of duckweed present, decreasing with increased duckweed coverage and at 50% duckweed coverage, their middle legs’ strides closely mimic their strides on sandpaper. Notably, *Microvelia* achieve speeds up to 56 body lengths per second on water, nearly double those observed on sandpaper and duckweed (both rough, frictional surfaces), highlighting their higher speeds on low friction surfaces such as the water’s surface. This study highlights *Microvelia*’s ecological adaptability, setting the stage for advancements in amphibious robotics that emulate their unique tripod gait for navigating complex terrains.

## Introduction

In nature, water surfaces are seldom clear; a pond’s surface is often littered with debris like fallen leaves, twigs from overhead trees, and small floating plants such as duckweed (family Lemnaceae), which can cover an entire pond’s surface [1]. This obstacle ridden surface is where neustonic insects such as water striders mainly traverse, contending with predators, competitors, and the challenge of walking on water [2, 3]. Studies have extensively explored the water strider’s locomotion mechanisms, which exploit surface tension through hydrophobic wax [4, 5] and dense hair coverage [6, 7]. However, research on the kinematics of transitioning across the varied substrates within their complex environments remains scant. Understanding these transitions has potential implications for future bio-inspired robots and morphological adaptations in real world settings [8, 9].

Of specific interest for substrate transition studies are *Microvelia*, water striders that uniquely navigate both water and land using a single gait: the alternating tripod gait (Fig. 1c) [10, 11]. Unlike them, other water walkers like Gerridae, which rely on a rowing gait, find land traversal challenging (if not impossible) due to their dependence on water contact for all legs [3]. Fishing spiders, Dolomedes, can move on both land and water but must switch gaits between the two [12, 13]. Some terrestrial, tropical ants adopt the alternating tripod gait for emergency water escapes but only for brief periods and with mixed success [14]. *C. schmitzi* ants, in symbiosis with pitcher plants, swim in digestive fluids but only for short durations and in limited capacity [15].

**Fig. 1.**
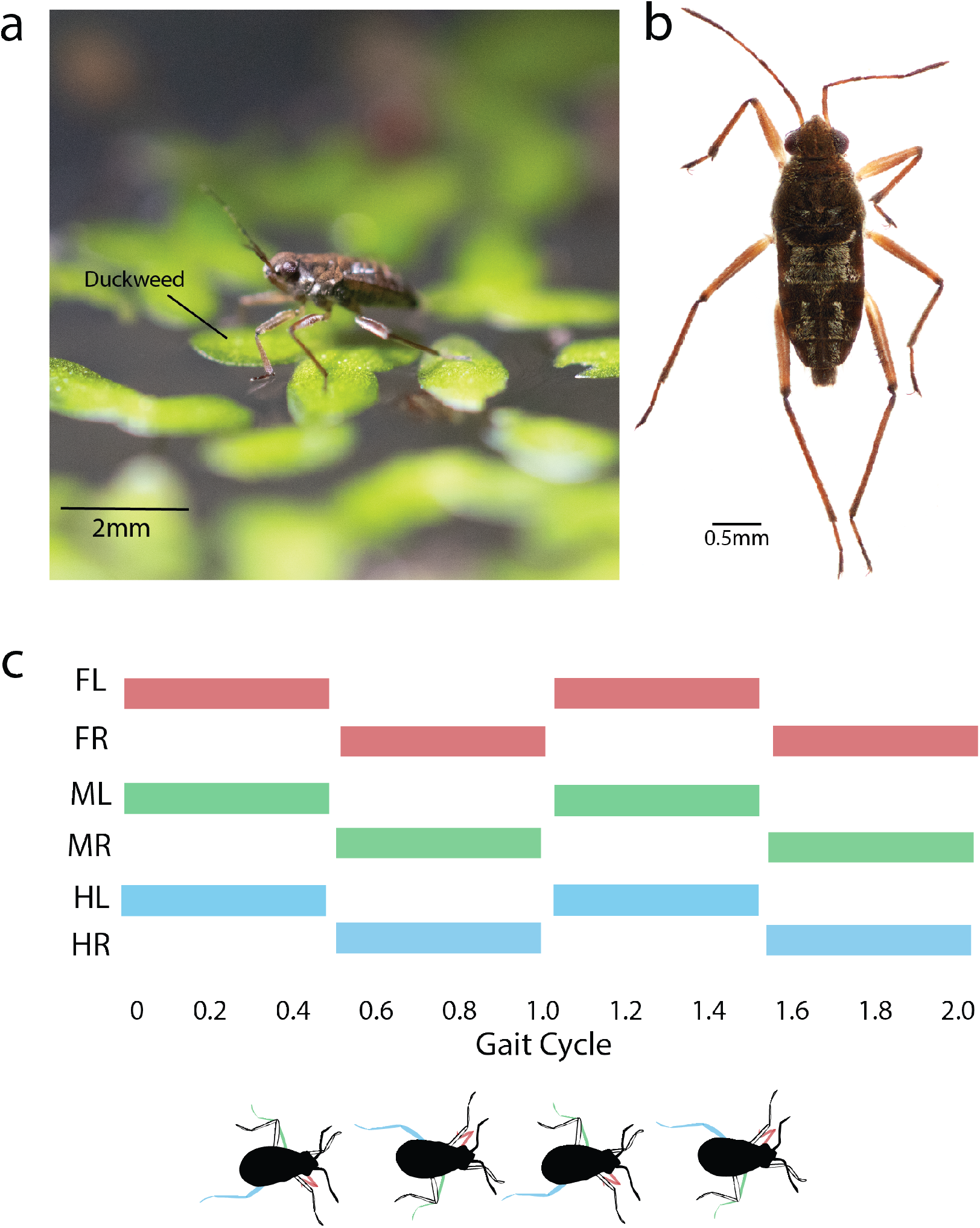
*Microvelia* and its alternating tripod gait. **(a)** A *Microvelia* standing on duckweed fronds. Image courtesy of Dr. Pankaj Rohilla. **(b)** High resolution image of a *Microvelia*. **(c)** Gait plot indicating the power stroke (colored rectangles) and recovery phase (blank rectangles) of the alternating tripod gait. The illustration below corresponds to the *Microvelia’s* gait cycle.

Robotics research has applied the alternating tripod gait on complex surfaces, primarily focusing on terrestrial environments [16]. Comprehensively understanding how *Microvelia* manages various substrates in its daily pond life, including rocks, debris, sand, water, and duckweed (Fig1a), is key for understanding the versatility of the alternating tripod gait that distinguishes it from its aquatic counterparts. This paper explores the multifaceted terrains *Microvelia* frequently navigates, offering new avenues for alternating tripod gait research. Understanding *Microvelia’s* consistent gait across different terrains opens potential for microrobots designed for robust travel across diverse landscapes [17, 18, 16].

We aim to examine *Microvelia*’s locomotive efficiency on water, duckweed-covered water, and dry sandpaper. Utilizing high-speed video and pose estimation software, we will analyze the kinematics – body speed, stroke amplitude, and frequency – of *Microvelia* as they navigate using the alternating tripod gait across these surfaces.

## Materials and methods

### Setup

We obtained *Microvelia* from ponds and creeks from Kennesaw, Georgia. The specimens were kept in a 17.5 X 14.0 X 6.5-inch^3^ plastic container. The container held water kept at a constant temperature of 20 °C and duckweed from the insects’ native bodies of water. The insects were provided with circadian lighting 12 hours out of the day, from 8 A.M. to 8 P.M. Additionally, the specimens were fed once each day with fruit flies procured from Carolina Biological Monday through Friday. We examined the locomotion of the specimens across three different surfaces: water, 1000-grit aluminum oxide sandpaper from Uxcell, and water covered with duckweed in varying degrees (10%, 25%, 50%). We estimated the duckweed percent coverage via binarizing an image of the surface with ImageJ [19]. In total, the locomotion of 3 specimens for each type of surface was examined.

### Recording

A Photron FASTCAM MINI AX 2000 set at a resolution of 1024 by 1024 pixels with a frame rate of 2,000 frames per second was used to record the locomotion of the *Microvelia* across the different surfaces. A Nikon 70-200mm f/2.8G ED VR II AF-S NIKKOR Zoom Lens was mounted onto the camera. The camera and lens were attached to a vertically placed Thorlabs Optical Rail and pointed at the specimens’ dorsal sides. We placed the insects into 10.0 by 10.0 by 1.5-centimeter Thermo Scientific petri dishes, each dish being either filled halfway with water, covered with 1000-grit sandpaper, or filled halfway with water and covered with varying amounts of duckweed (Fig2). Each insect was recorded individually and prodded for movement. These petri dishes were raised slightly above a table, put against a white background, placed directly under the camera’s lens. An LED light was also lit underneath the petri dishes for enhanced recording quality.

**Fig. 2.**
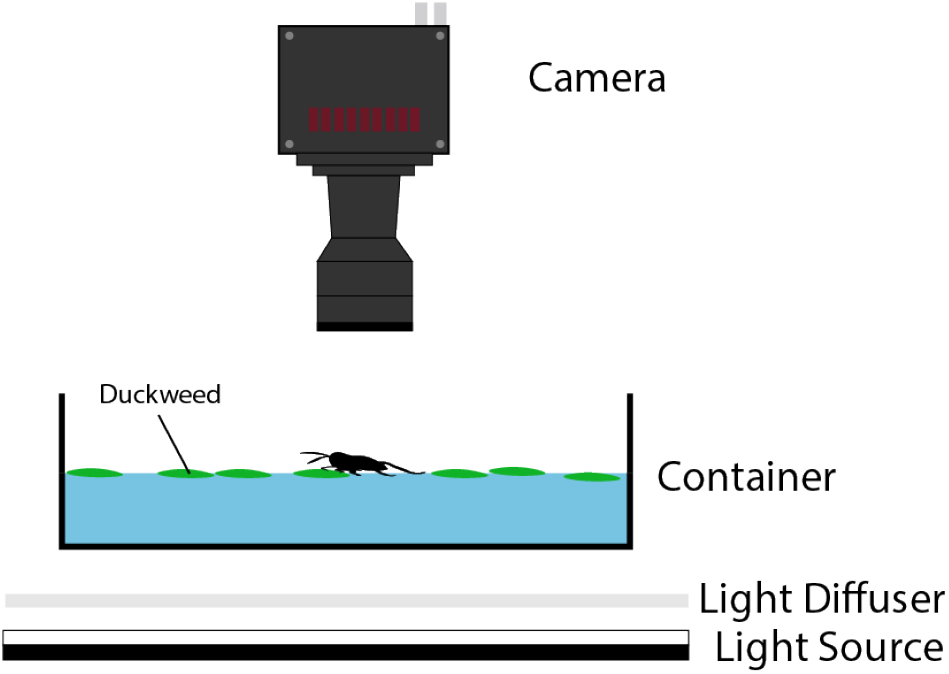
Experimental Setup. Schematic of experimental setup. A high speed camera is mounted above a container of water with duckweed on top. The container rests on a diffuser. A light source is set at a short distance below the diffuser to provide more even lighting. *Microvelia* are recorded individually running on the water partially covered with duckweed.

### Tracking, Data Acquisition, and Analysis

After recording, the following points on the specimen were tracked for each recording: the coxae, tibiofemoral joints, tibiotarsal joints, tarsi tips, the abdomen tip, and the head. DeepLabCut (DLC) pose estimation machine learning software was utilized entirely for the sandpaper and water surfaces when it came to tracking of all the aforementioned points [20] (see Supplementary Movies S1 and S2). However, on the water-duckweed surface, DLC was used only to track the tip of the abdomen, the head, the coxae, and the tibiofemoral joints. We tracked the rest of the points manually using PFV4 since DLC was unable to track these particular points with sufficient accuracy (see Supplementary Movie S3). Ultimately, we used the data gathered from the videos and the trackings (position and time of each point) to calculate the displacement, velocity, joint angles, and step amplitude for each recorded specimen. The kinematics of the left and right leg for each pair were averaged together. For statistical analysis we used one-way ANOVA test to determine if there was a difference amongst the means of each group with Tukey’s difference criterion to find which pairs were statistically different.

## Results

### Kinematics of Running on Duckweed, Sandpaper, and Water

*Microvelia* (Fig1b) traverse both land and water, though prior research primarily focused on smooth substrates, neglecting plant-surface substrates (e.g. duckweed Fig1a), and lacked within-species comparisons [3]. To address this, we tested *Microvelia* on high friction sandpaper to mimic the rough terrain (rocks) surrounding their aquatic habitats and on duckweed-covered water surfaces to assess locomotion on natural, heterogeneous surfaces within their environment. Across all substrate types – frictionless water, high friction sandpaper, and water with intermittent friction from duckweed coverage – *Microvelia* exhibited the alternating tripod gait (Fig1c). This gait is widespread amongst hexapods on land [21] and is adapted by swimming ants in a modified form [14, 15]. Along with noticeable visual differences in the gait (Fig4a), we identified distinctive gait properties for *Microvelia* across the three different substrates, described in the next sections.

#### Body and leg velocity

We found *Microvelia* is significantly faster on water, achieving a maximum speed of 56 body lengths per second (bl/s) (Fig3a, *p <* 0.001). In contrast, its maximum body speeds on sandpaper and with 50% duckweed coverage, at 26.5 bl/s and 28.7 bl/s respectively (*p >* 0.05), are about half that on water. Across substrates, *Microvelia*’s upper legs move at similar maximum speeds (Fig3b). Yet, on water, *Microvelia’s* middle and hind legs moved faster than on sandpaper and duckweed at 51 bl/s (*p <* 0.001) and 46 bl/s (*p <* 0.001) respectively. This trend mirrors the body speed observations, which might explain the lower body speeds on sandpaper and 50% duckweed, where the middle and hind legs did not exceed speeds of 40 cm/s.

**Fig. 3.**
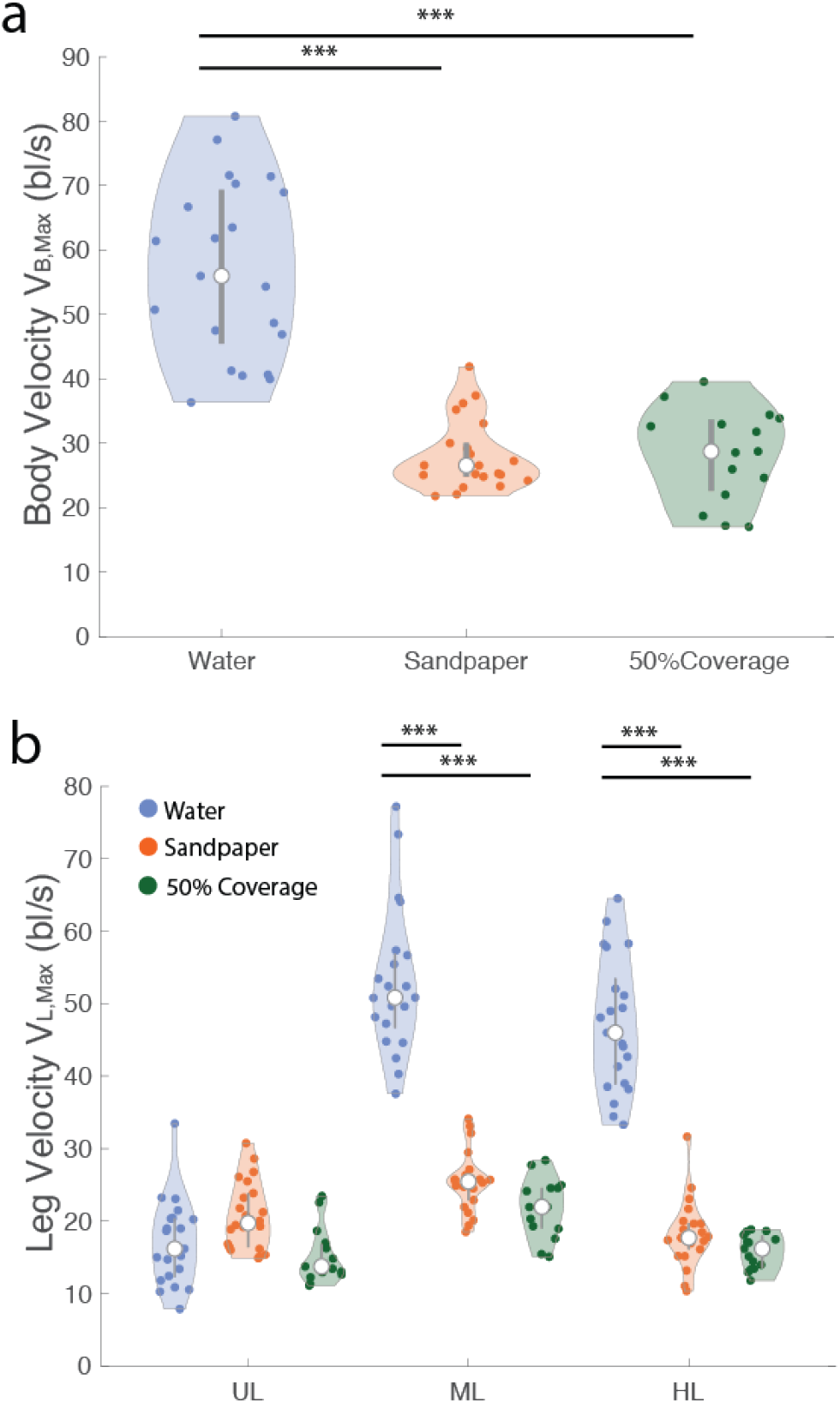
Maximum velocities of body and of legs across substrates. Maximum velocities *V*_max_ of *Microvelia* on sandpaper and duckweed are comparable, whereas movement on purely water is distinct. **(a)** Body velocity comparison on water, sandpaper, and 50% duckweed coverage. **(b)** Leg velocity comparison on water, sandpaper, and 50% duckweed coverage for each leg location (upper legs, middle legs, and hind legs). White circles represent the median. Bar represents 2nd and 3rd quartiles. p-values: * *p <* 0.05, ** *p <* 0.01, *** *p <* 0.001.

**Fig. 4.**
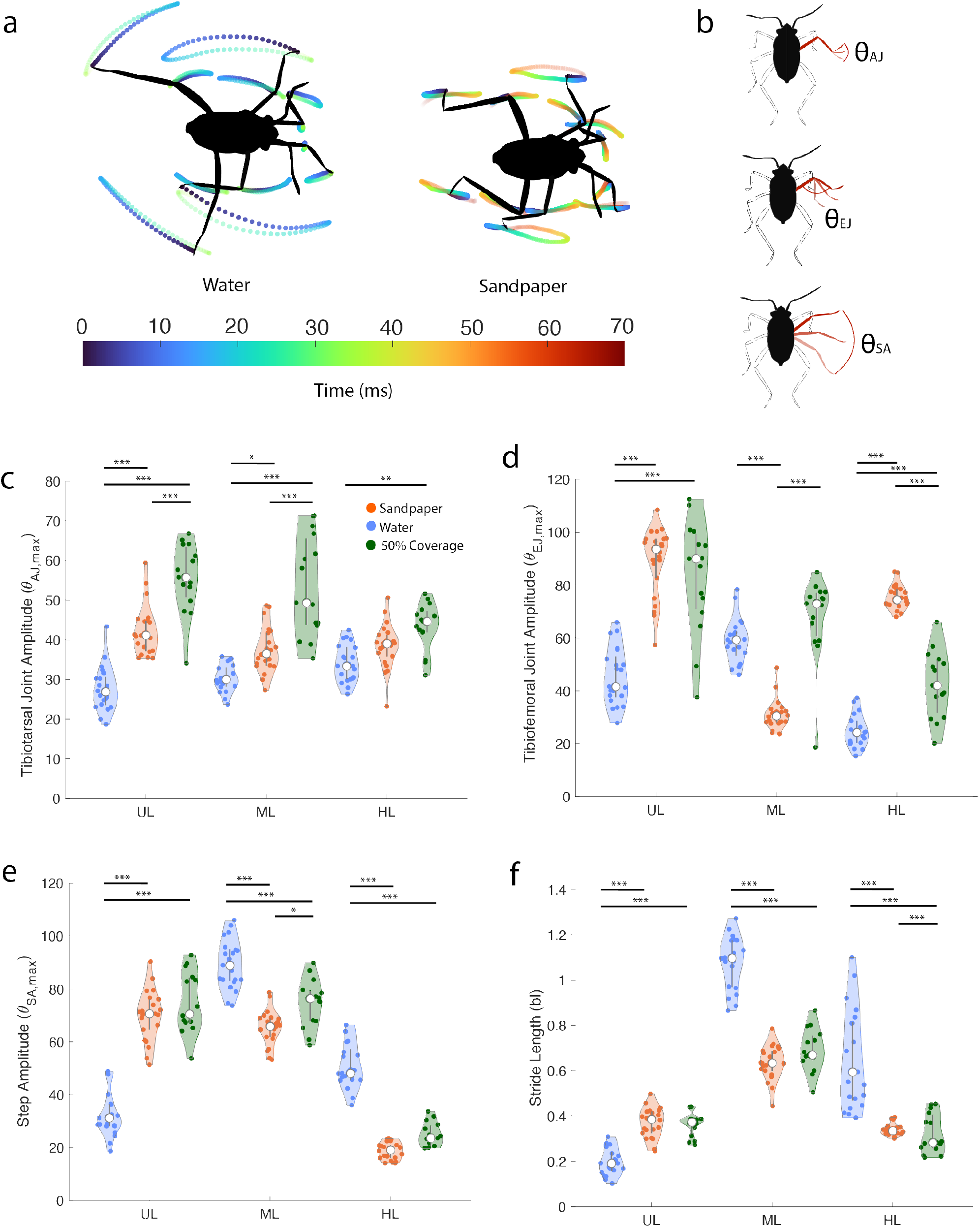
Kinematics of *Microvelia* locomotion on different substrates. **(a)** *Microvelia* tarsi and tibiofemoral joint trajectories on sandpaper compared to on water **(b)** Schematic demonstrating how each angle is calculated. From top to bottom: tibiotarsal joint (AJ), tibiofemoral joint (EJ), and stroke amplitude (SA). **(c-e)** Amplitudes of each leg, upper leg (UL), middle leg (ML), and hind leg (HL), according to the joint angles illustrated in (b), across substrates. **(f)** Stride length comparison across substrates. White circles represent the median. Bar represents 2nd and 3rd quartiles. p-values: * *p <* 0.05, ** *p <* 0.01, *** *p <* 0.001.

#### Joint angles

We measured the tibiotarsal joints and tibiofemoral joints, along with step amplitudes for all legs across the three different substrates (4b). The ampltiude of the tibiotarsal joints (*θ*_AJ,max_) showed an increasing trend from water to sandpaper to duckweed for all legs (*p <* 0.05*a*, Fig4c). On both duckweed and sandpaper, the amplitudes of tibiofemoral joints (*θ*_*EJ, max*_) for the upper and hind legs were higher than those on water (*p <* 0.001). The middle leg presented an exception, as its *θ*_*EJ,max*_ was lowest on sandpaper (*p <* 0.001). In terms of tibiofemoral joints, both upper leg and hind legs exhibited higher amplitudes on sandpaper and duckweed compared to water (*p <* 0.001).

#### Step amplitudes and stride lengths

For water, the step amplitudes and stride lengths are lowest in the upper legs and highest in the middle and hind legs (Fig4e-f, *p <* 0.001, see Supplementary Movie S4). Fig4a illustrates the increase in stride length for water compared to sandpaper. The lower step amplitudes and stride lengths in the upper legs are consistent with prior studies indicating that upper legs serve as stabilizers, having limited movement for water striders on water [3]. The middle leg, identified as *Microvelia’s* main propulsor on water, similar to other water striders [22], showcases relatively high stride lengths on water (1.1 bl, Fig4f). Interestingly, the step amplitudes and stride lengths for both middle leg and hind legs decrease in the presence of solid surfaces (duckweed and sandpaper, see Supplementary Movies S7 and S8), correlating with their reduced maximum velocities on these higher friction surfaces (Fig3). On water, while the hind leg’s tibiofemoral amplitude remains low, its stride lengths are higher, whereas on duckweed, despite shorter stride lengths than on water, the tibiofemoral amplitude increases. For the upper legs, an increase in stride length accompanies rising tibiofemoral amplitude.

### The Effect of Duckweed Coverage on Stride Length

We compare the average stride lengths of the upper, middle, and hind legs across water, sandpaper, and three levels of duckweed coverage (10%, 25%, and 50%, Fig5a-c). We found no statistical difference in stride lengths among all duckweed coverages and sandpaper (*p >* 0.05), indicating that *Microvelia* exhibits similar stepping behavior on duckweed and sandpaper, regardless of surface coverage by obstacles. The stride lengths of both upper and hind legs show that *Microvelia* approaches all levels of solid substrates with a uniform stepping pattern.

**Fig. 5.**
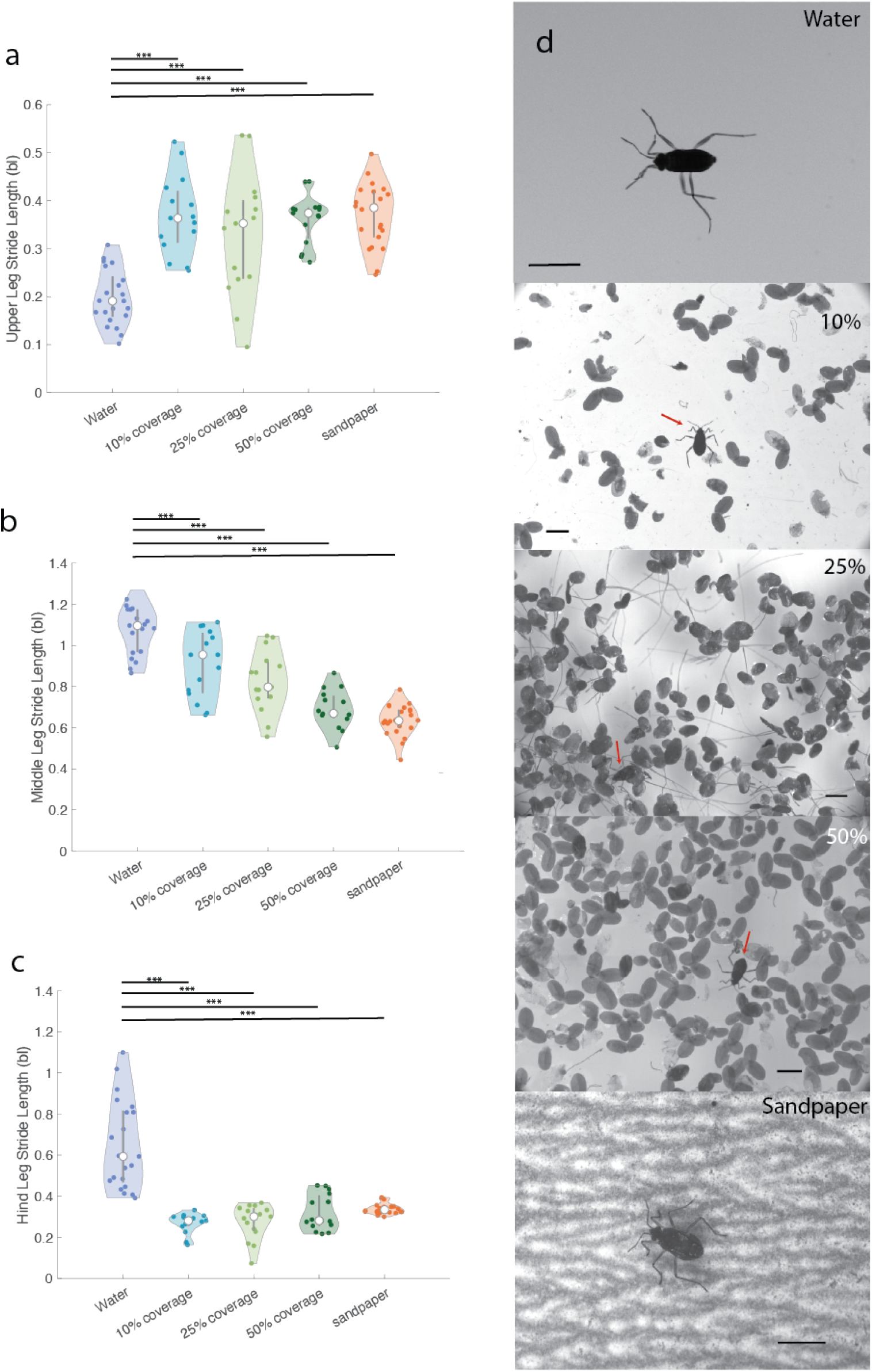
Stride length comparison across substrates and varied duckweed coverages (10%, 25%, 50%). **(a)**Average stride length of upper legs on each substrate shows that *Microvelia* increases their upper leg’s stride at the presence of a solid substrate. **(b)** Average stride length of middle legs across substrates show that increase in duckweed coverage leads to a decrease in stride length. At 50% coverage the stride length of the middle leg is similar to the stride length on sandpaper. **(c)** Average stride length of hind legs reveals that *Microvelia* decrease the stride of their hind lengths at the presence of a solid substrate. **(d)** Photos of each substrate with an individual *Microvelia*. Each scale bar represents 2mm. Red arrows shows where *Microvelia* is located. From top to bottom, the substrates are clear water, water with 10% duckweed coverage, 25% duckweed coverage, 50% duckweed coverage, then sandpaper. Duckweed is sometimes found with submerged routes underneath the frond as seen in 25% coverage image. White circles represent the median. Bar represents 2nd and 3rd quartiles. p-values: * *p <* 0.05, ** *p <* 0.01, *** *p <* 0.001.

Mirroring trends identified in previous studies on water, we noted that the upper and hind legs display smaller step amplitudes and stride lengths, with the upper legs serving as ‘stabilizers’ and hind legs functioning as ‘rudders’ [3]. The stride length of the upper legs is found to be the shortest on water (0.19 bl, N=3, n=21) compared to other substrates (Fig5a, *p <* 0.001). In contrast, the stride lengths of the middle and hind legs are higher on water than on duckweed and sandpaper (Fig5b-c, *p <* 0.001), showcasing an inverse trend. Specifically, the stride lengths of the middle legs decrease as the friction or heterogeneity of substrates (% duckweed) increases (Fig5b, *p <* 0.001). With *Microvelia* moving slower on solid substrates (Fig3a-b), their upper legs became more active, displaying higher joint angles (Fig4).

## Discussion

Navigating complex environments necessitates that organisms adapt or modify their gait for survival. Hexapods, such as cockroaches and ants, employ the alternating tripod gait and modify it for various purposes. Blaberid cockroaches switch from the alternating tripod gait to a metachronal gait, reducing vertical amplitudes and enhancing lateral amplitudes to speed up on land [23]. Similarly, wood ants [24] and fruit flies [25] increase their stride frequencies to hasten land movement. North African desert ants shorten their stance phase to boost their body speed [26]. These adaptations primarily aim to increase terrestrial body speed and do not apply to aquatic environments.

Focusing on the air-water interface reveals different gait changes and adaptations for water locomotion. Tree canopy ants, *Pachycondyla spp*. and *O. bauri*, use their contralateral front legs and middle legs to row across water surfaces in a modified alternating tripod gait [15], using their hind legs for roll stability to prevent from flipping over. Conversely, tree canopy ants *C. americanus*, use their middle legs as rudders rather than for rowing [15]. These adaptations serve temporary excursions into fluids, with *C. schmitzi* ants, for instance, which live symbiotically with the pitcher plant, staying in fluid for less than 45 seconds [14]. Other water-dwelling organisms, like the fisher spider, switch from a rowing gait on water to an alternating tetrapod gait on land [27, 13]. Unlike these examples, water striders such as Gerridae, Rhagovelia, and Velia rely on a rowing gait and do not transition this gait to terrestrial locomotion [22, 3, 11, 28, 29, 22].

For the tiny *Microvelia*, navigating a pond’s complex and obstacle-laden environment necessitates adaptability over all surfaces. Our findings reveal that *Microvelia* not only locomotes on water, duckweed, and sandpaper but also adapts to the variation of these surfaces.

### Upper legs and hind legs contribute more on land vs. water

*Microvelia* achieve significantly higher speeds on water than on land or duckweed-covered areas (Fig3), as demonstrated by their increased step amplitudes and speeds on water (Fig4e). The middle legs display longer stride lengths and larger step amplitudes than the other legs when on water (see Supplementary Movie S4), consistent with previous studies that assign the role of stability to the upper legs, propulsion to the middle legs, and rudder function to the hind legs [3]. Acting as oars, the water strider legs push against the water [30, 31], with the middle legs stroking at a higher amplitude to provide the most propulsion. This action suggests that decreasing the tibiofemoral joint amplitude in the hind legs could lead to less power use, more energy conservation while pushing against the frictionless smooth surface of water [32]. On sandpaper and with 50% duckweed coverage, however, *Microvelia* increase their hind legs’ joint angles while reducing their stride lengths and step amplitudes (see Supplementary Movies S7 and S8). They also heighten the joint angles in their upper legs along with increasing stride lengths and step amplitudes (Fig4c-f). This adjustment occurs because *Microvelia* bend their legs more, possibly lifting them higher to navigate the topology of frictional rough surfaces. On such surfaces, *Microvelia* face difficulty sweeping and extending their legs as easily as on water, due to obstacles obstructing their tarsi, leading to shorter stride lengths and greater leg bending in the upper and hind legs. Foot trajectory comparisons on frictionless water versus frictional land surfaces further illustrate these differences (Fig4a). Terrestrial insects using the alternating tripod gait, like cockroaches, also show higher hind tibiofemoral joint amplitudes on frictional surfaces [21, 33].

### Microvelia treat duckweed as a land-like surface

Across all substrates, *Microvelia’s* middle legs demonstrate the least variance in stride lengths, especially when comparing water, various duckweed coverages, and sandpaper (Fig5 a-c). These legs also maintain tibiofemoral joint angle values relatively consistent (Fig4d). This consistency suggests that the middle legs, known for being the main propulsors on water, maintain a similar function across different terrains. Our findings indicate that *Microvelia* navigate duckweed coverages similarly to how they would navigate sandpaper, treating both as “land” conditions. While adapting their gait to accommodate substrates floating on water, *Microvelia* distribute more work to the other legs on land-like surfaces as the middle leg stride lengths decrease on duckweed and sandpaper.

This unique application of a common terrestrial gait, the alternating tripod gait, for aquatic running showcases the potential for bioinspired designs in cross-terrain and amphibious micro-robots. These adaptations highlight opportunities for further research in gaits adjustments across substrates, the biological actuation behind traversal in diverse environments, and the implications for semi-aquatic robotics and bio-inspired design in navigating complex media such as sand [34, 16]. Inspired by *Microvelia*, future robotic designs might only require a single adaptable gait for multifaceted environmental navigation, offering insights into mechanosensory affordances for multi-environmental adaptability [27, 16, 18, 35].

### Limitations

While aiming to mimic the complex and varied system of a pond during our *Microvelia* recordings, the range limitations of our high-speed camera and ability to reliably use DeepLabCut to track joints in this tiny insect constrained the area available for *Microvelia* locomotion. Our study also encompassed a small sample size and examined a preliminary selection of substrates (limited % of duckweed coverage and 1 sandpaper type). Despite these constraints, our experimental setup yielded consistent results across tests. Future studies could expand the number of specimens, possibly including different juvenile instars for developmental comparisons and explore additional substrates or duckweed coverage densities. We observed a z-component in the amplitude of leg and joint movements, which our study did not capture. Accurately tracking leg movements in the z-direction would offer a more complete understanding of leg behavior on heterogeneous, rough surfaces.

We also noted that duckweed fronds move when *Microvelia* traverse them. Future studies could quantify the movement of these fronds during *Microvelia* tarsi interactions. Examining locomotion on wet versus dry surfaces could provide additional insights, given that *Microvelia* inhabit environments where they may encounter both (after a rainshower). *Microvelia’s* primary movements – to pursue prey or escape predators (the latter being the focus of this study) – mean that their cross-substrate locomotion is not always continuous. For instance, on surfaces with sparsely scattered duckweed, *Microvelia* often move across larger water areas and halt upon reaching duckweed. This behaviour likely serves as a underwater-predator evasion strategy, yet it limited our observations of smooth transitions between aquatic and terrestrial locomotion.

## Conclusions

In our study, we determined how *Microvelia* modifies the alternating tripod gait to traverse across different surfaces through high speed imaging and pose-estimation deep-learning software. Through our results, we discover that *Microvelia* move their upper legs at a higher stride length on land than water, suggesting that the upper legs provide more propulsion on land and may be needed to facilitate walking on rougher terrain. Furthermore, we discover that the stride lengths of the upper legs and hind legs are statistically similar across all duckweed coverages and sandpaper within this study. This suggests that once *Microvelia* know that solid debris is present on water, that they will adjust their upper and hind legs to move similarly to their movement on land. *Microvelia* were also found to decrease their step amplitude with increasing duckweed coverage. Since the middle legs are used as the main means of propulsion, our data suggests the *Microvelia* are adjusting the stride of their middle legs to move more quickly on more variable terrain. Ultimately, these results can influence the design of future amphibious microbots that can better traverse rough and uncertain terrain that may include random debris.

## Supplementary data

Supplementary data available at *ICB* online.

## Data Availability

The data underlying this article are available in the article and in its online supplementary material.

## Author contributions statement

J.O and H.W designed the experiments. J.O, K.Y, G.D, and H.W. carried out the experiments and acquired the data. J.O analyzed the data and interpreted the results. M.S.B. reviewed the design and execution of experiments, the data analysis, the interpretations, and the manuscript. J.O, K.Y, G.D, and M.S.B contributed to writing the manuscript.

## Acknowledgments

J.O. acknowledges funding support from the GT UCEM fellowshp program and the Herbert P. Haley fellowship program. M.S.B. acknowledges funding support from the NIH Grant R35GM142588 and NSF Grants CAREER 1941933 and 2310691. We thank members of the Bhamla lab for useful discussions and feedback of this work and Dr. Pankaj Rohilla for feedback and mentorship.

