## Supplementary material for "Tiny amphibious insects use tripod gait for seamless transition across land, water, and duckweed": Figures SI_ICB.pdf

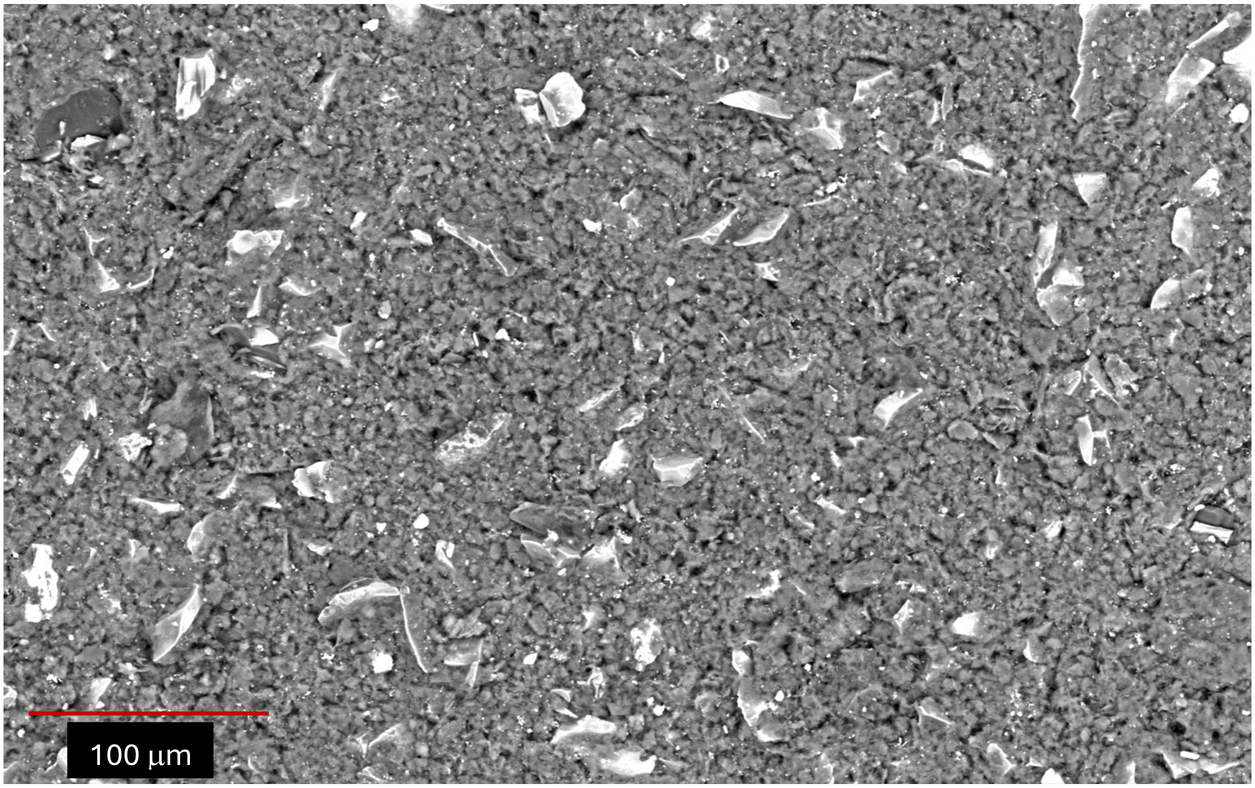


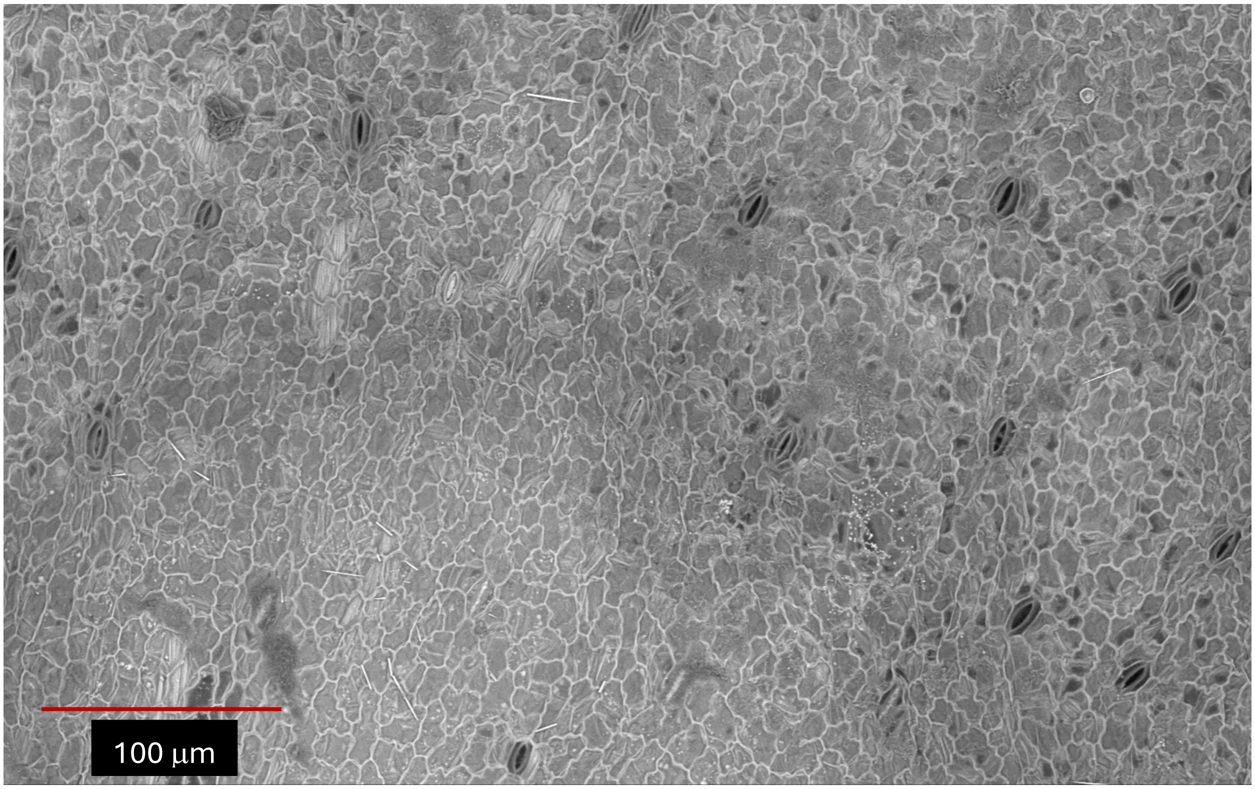


**Fig S1:** Scanning Electron Microscopy Images of 1000 grit sandpaper (top) and duckweed (bottom).

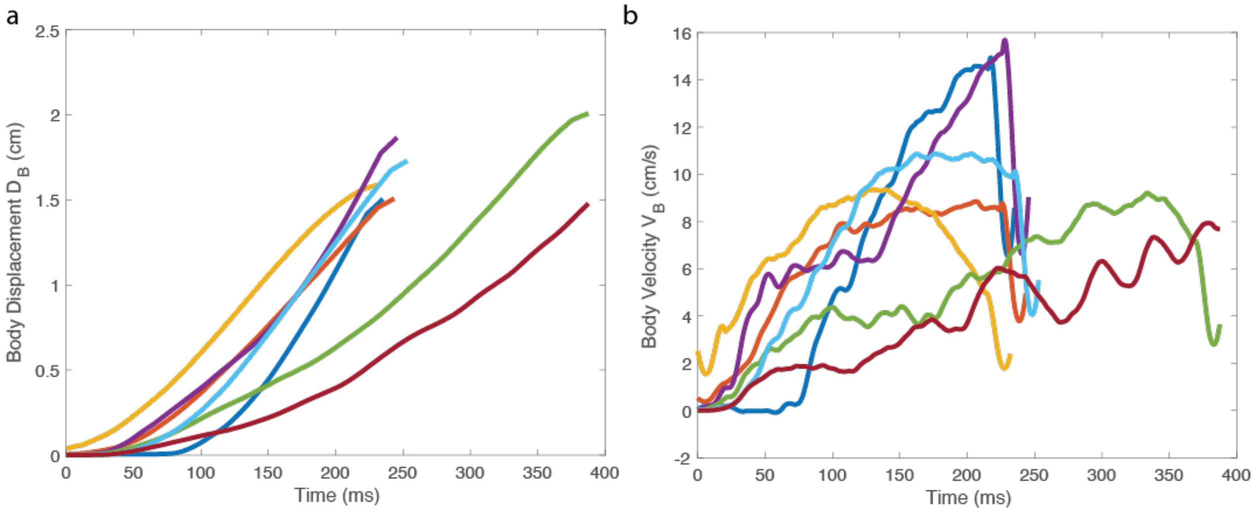


**Figure S2: Body Displacement and Body velocity over time for a Non-Ablated *Microvelia*.** (a) Displacement versus time of a single Microvelia. (N=1 specimen, n=7 trials) (b) Velocity versus time of a single Microvelia. (N=1 specimen, n=7 trials)

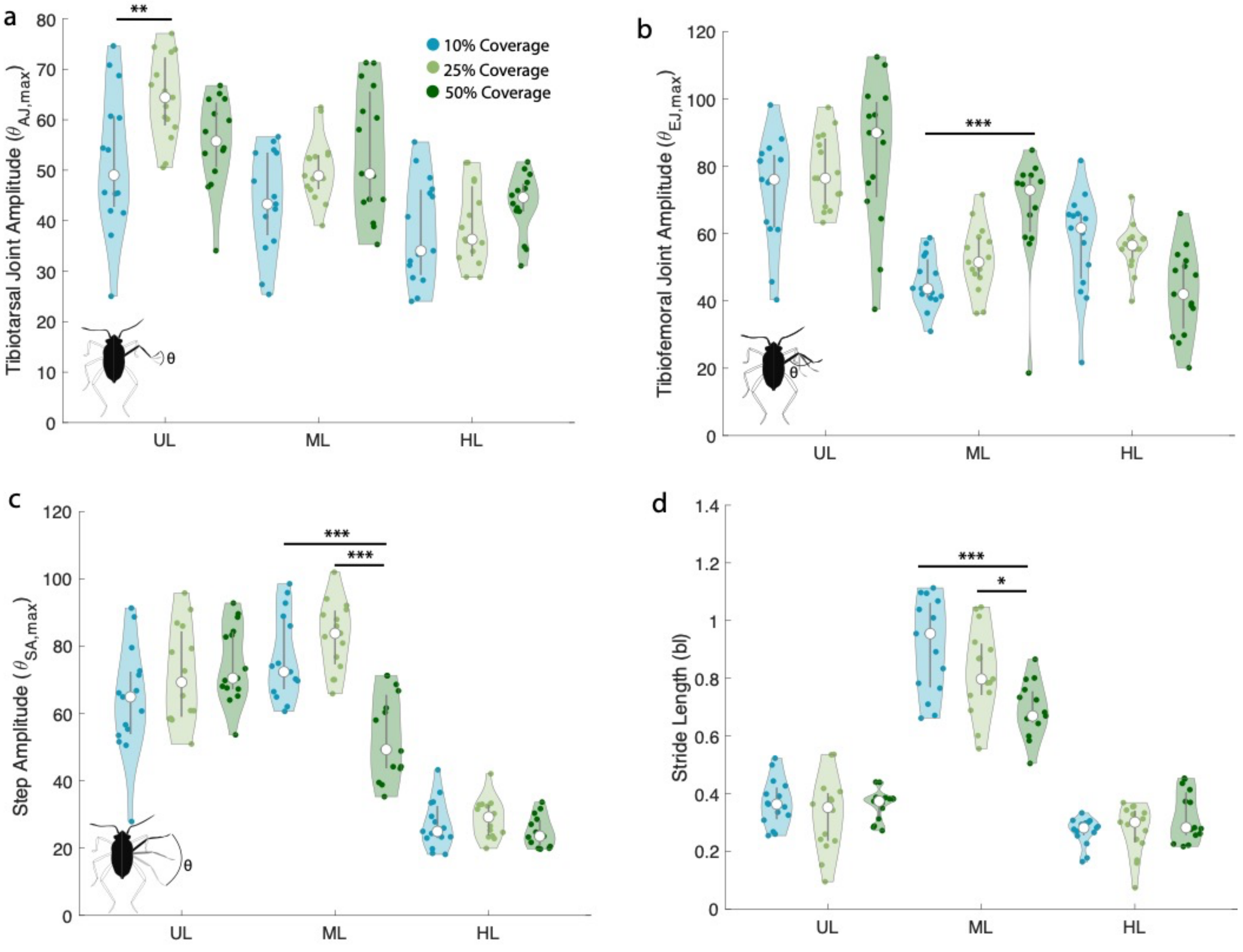

**Figure S3: Kinematics of *Microvelia* locomotion on different duckweed coverages**. **(c-e)** Amplitudes of upper leg (UL), middle leg (ML), and hind leg (HL), according to the joint angles illustrated across duckweed coverages. **(f)** Stride length comparison of each leg across substrates. The kinematics of the left and right leg in c-f for each pair were averaged together. P-values: * P<0.05, ** P<0.01, *** P<0.001.

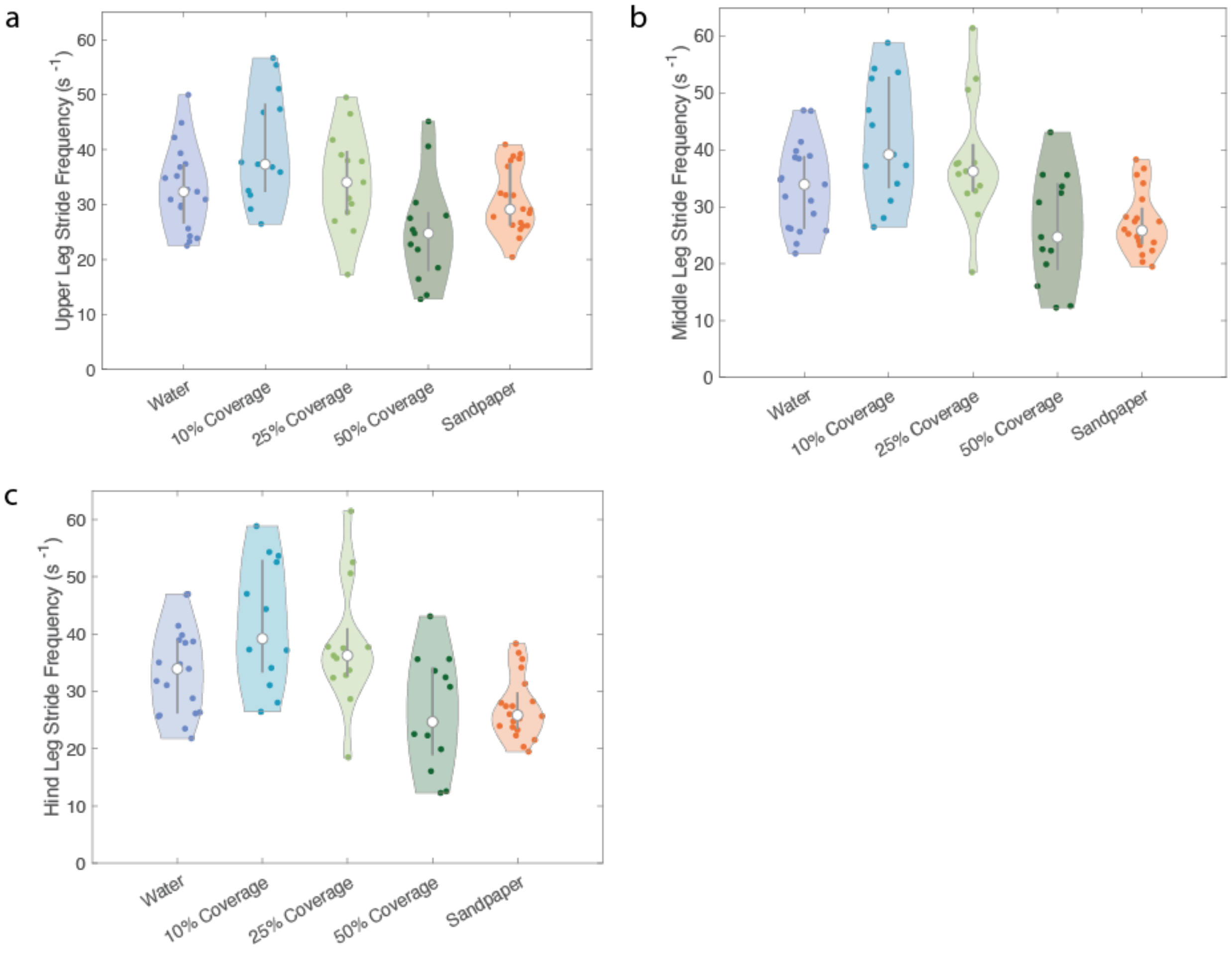


**Figure S4: Stride Frequency of Each leg**. *Microvelia* Stride frequency for (a) upper, (b) middle, and (c) hind legs.

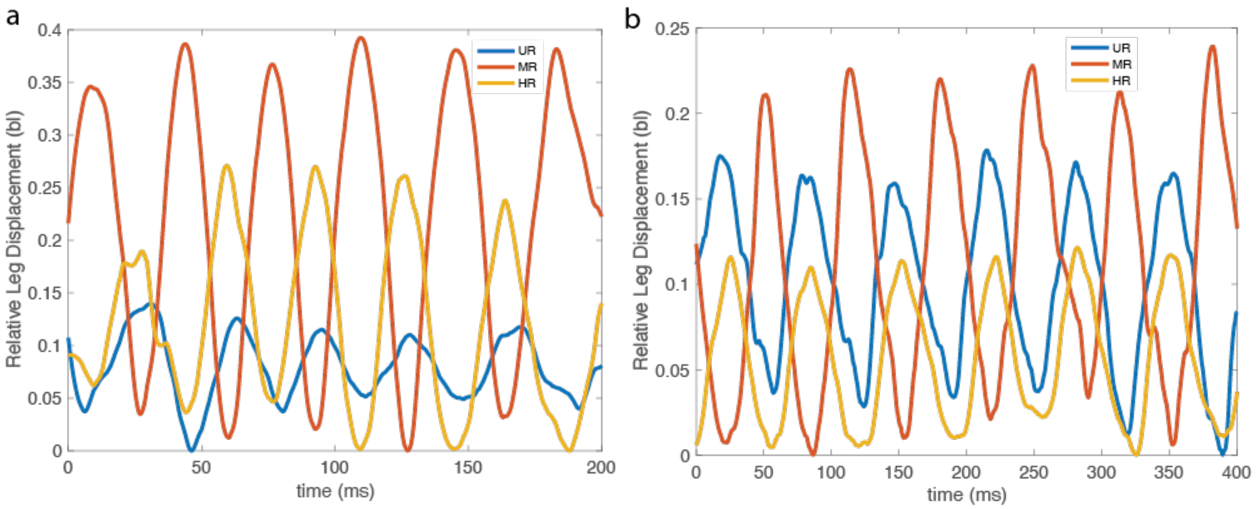


**Figure S5: Relative Leg Displacement:** Relative Leg Displacement on water (a) and sandpaper (b) for all right legs. The displacement is relative to the lowest point of the tarsal tip relative to the coxa during stance phase. Stride length is the peak height of each peak.
